# Association between hot flashes severity and oxidative stress among Mexican postmenopausal women: a cross-sectional study

**DOI:** 10.1101/575621

**Authors:** Martha A. Sánchez-Rodríguez, Mariano Zacarías-Flores, Alicia Arronte-Rosales, Víctor Manuel Mendoza-Núñez

**Affiliations:** Unidad de Investigación en Gerontología, Facultad de Estudios Superiores Zaragoza, Universidad Nacional Autónoma de México, Ciudad de México, México; División de Ginecología y Obstetricia, Hospital Gustavo Baz Prada, Instituto de Salud del Estado de México, Nezahualcoyotl, Estado de México, México

**Keywords:** oxidative stress, postmenopausal women, hot flashes, malondialdehyde, antioxidant status

## Abstract

**Objective:** To assess the association between hot flashes (HFs) severity and oxidative stress (OS) in Mexican postmenopausal women.

**Methods:** A cross-sectional study was carried out with perimenopausal women aged 40-59 years community-dwelling from Mexico City, Mexico. They participated in Menopause and Oxidative Stress Project. The baseline sample consisted of 476 women recruited to participate; 161 women were excluded due to different reasons. Hence, 315 women were selected to establish two groups, a) 145 premenopausal women (yet with menstrual bleeding), and b) 170 postmenopausal women (without menses). All women were free of cardiovascular, kidney, hepatic or cancer disease, and without antioxidant supplement intake for at least six months prior to the beginning of the study; none had previously received hormone therapy. As OS markers, we measured plasma malondialdehyde using the TBARS assay, erythrocyte superoxide dismutase (SOD) and glutathione peroxidase (GPx), uric acid, and total antioxidant status; also, we calculated SOD/GPx ratio, antioxidant gap and an oxidative stress score ranging from 0 to 7. The HFs were evaluated using the Menopause Rating Scale. The women completed Spanish version of the Athens Insomnia Scale, Zung Self-Rating Anxiety Scale and Zung Self-Rating Depression Scale and a questionnaire of pro-oxidant factors.

**Results:** Stress score increased with HFs severity (mild 2.9±0.23, moderate 3.1±0.21 and severe 3.8±0.18, p<0.01) in postmenopausal women. We observed a positive correlation between HFs severity and stress score, r=0.247 (p=0.001) in postmenopausal women; other test scores were not correlated. Severe HFs were a risk factor for OS (OR=3.37, 95%CI: 1.20-9.51, p<0.05) in an adjusted multivariate analysis by different postmenopausal symptoms and pro-oxidant factors; we did not see any association in premenopausal women.

**Conclusion:** Our findings suggest an association between HFs severity and OS in Mexican postmenopausal women.

## Introduction

Menopause, an expected event in a woman’s life, is commonly defined as a 12-month period of amenorrhea [1] or hypoestrogenism (estrogen level < 25 pg/mL) due to ovarian senescence; therefore, the postmenopausal period may be considered the beginning of the aging process in women. Postmenopausal aging is produced by a series of endocrinological changes that lead to the erratic production of estrogens (mainly estradiol) that eventually bring to low estrogen (E2) level [2] and is associated with multiple symptoms including vasomotor symptoms that interfere with daily activities and sleep. The most distressing symptoms of menopausal transition are hot flashes (HFs). They occur in over 75% of menopausal women [3]. Recently, it was highlighted that moderate/severe HFs continue, on average, for nearly 5 years after menopause, and more than one third of women experience moderate/severe HFs 10 years or more after menopause [4]; however, HFs onset or intensify occurs during the late menopausal transition [1].

The marked reduction in E2 has been shown to increase levels of oxidative stress (OS) in the body because E2 presents antioxidant properties due to its structure and its capacity to prevent OS by different ways, such as free-radical scavenger, neutralizing excess reactive oxygen species (ROS), and increasing antioxidant molecules (e.g. thioredoxin and superoxide dismutase) [5,6]; therefore, E2 is part of the antioxidant system that counteracts OS during reproductive stage. Additionally, low concentrations of this hormone have pro-oxidant like effects [7]. In this regard, our research group have described that menopause is a risk factor for OS [8] because when the production of E2 decreases, the antioxidant protection is lost and therefore OS increases.

Oxidative stress is also involved in the pathogenesis of menopausal symptoms, such as vasomotor disturbances (e.g. HFs or night sweats). During menopause transition and postmenopausal period, the women suffer repeated episodes of such vasomotor disturbances, which produce an increase of the metabolic rate. These episodes of vasomotor symptoms contribute to OS production by raising the level of oxidant species and by blocking antioxidants and their function in neutralizing reactive oxygen/nitrogen species [7].

Additionally, the relationship between HFs and OS is little understood; several studies support an association, but others do not. Recently a report noted that HFs and OS are independent events [9], causing a controversy; therefore, the aim of this study was to assess the association between HFs severity and OS in in Mexican postmenopausal women.

## Material and methods

### Study design and population

We carried out a cross-sectional study with a deterministic sample of 315 perimenopausal women aged 40-59 years community-dwelling from Mexico City, Mexico. They were invited to participate in Menopause and Oxidative Stress Project directed by the Gerontology Research Unit at Universidad Nacional Autonoma de Mexico, Zaragoza Campus, from February 2015 to March 2016. The baseline sample consists of 476 women recruited by informative brochures that were distributed in the community specifying the objectives of the study and the admission criteria; 161 women were excluded due to different reasons (Fig 1). Women included were separated into two groups, a) 145 premenopausal women (yet with menstrual bleeding), and b) 170 postmenopausal women (without menses). All women were free of overt cardiovascular, kidney, hepatic and cancer disease as assessed by medical history and physical examination and without antioxidant supplement intake for at least six months prior to the beginning of the study; none had previously received hormone therapy. The study protocol was approved by the Ethics Committee of the Universidad Nacional Autonoma de Mexico, Zaragoza Campus, Mexico City, Mexico (register number FESZ/DEPI/CI/004/17).

**Fig 1.**
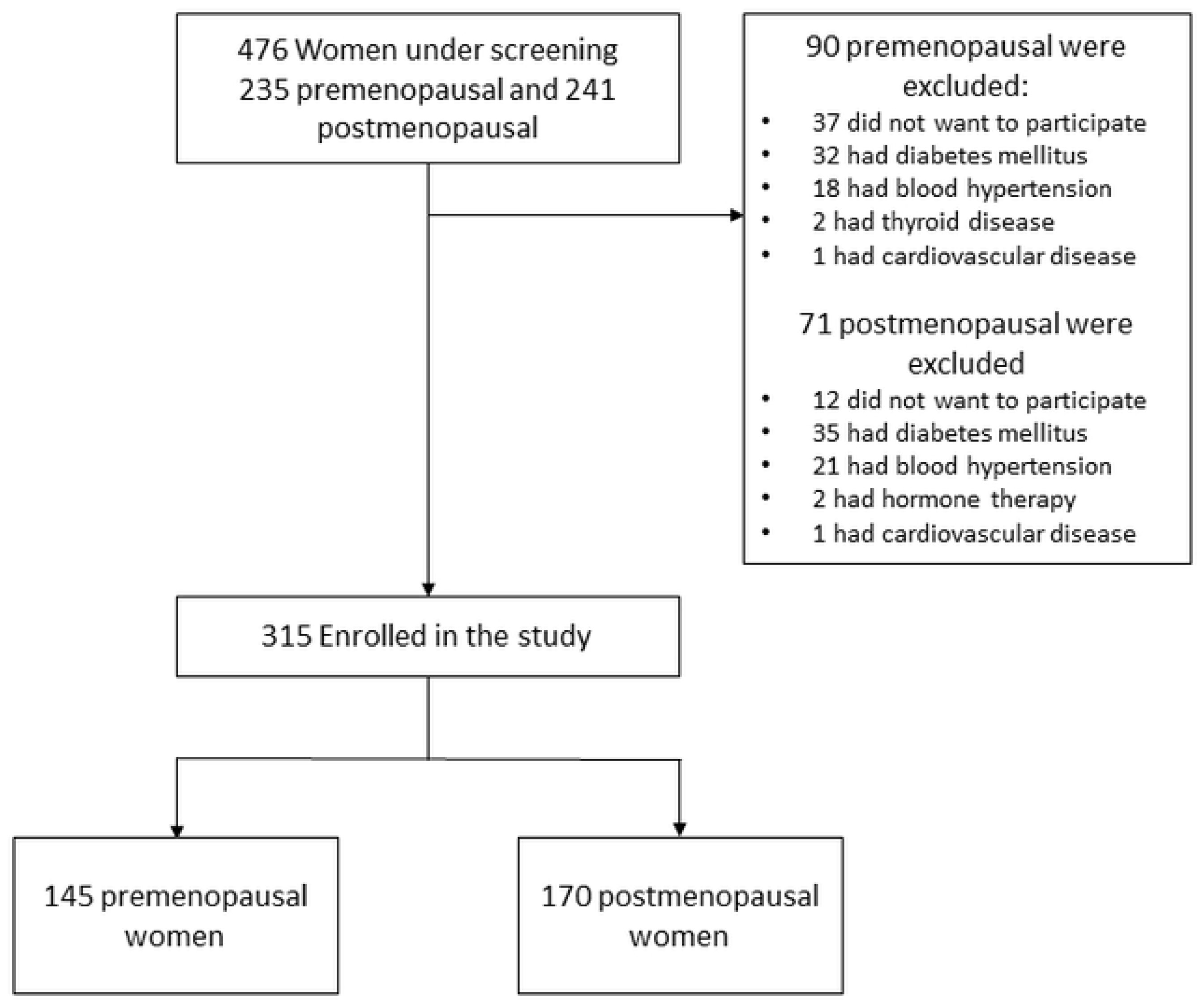
Diagram of the inclusion of study participants.

The women agreed to participate in the study after signing the informed consent. The participants underwent the following examinations: complete clinical history, complete blood count, glucose and lipids profile, anthropometric and blood pressure measurements. Those tests were used to establish their health status, using the cut-off points of reference values for Mexican adults [10].

We measured E2 level using a radioimmunoassay method (Siemens, Malvern, PA, USA) and FSH level using a chemiluminescence method (Siemens). The within-run precision level for these methods were 3.1% and 7.4%, respectively, and the E2 analytical sensitivity was 8 pg/mL.

Blood samples were collected after a 12-h fasting period by venipuncture and placed in vacutainer/siliconized test tubes containing a separating gel and no additives, and heparin as anticoagulant agent (Becton-Dickinson, Mexico City, Mexico). Samples containing heparin were analyzed using a hemoglobin test protocol (including hemoglobin, hematocrit, and leukocyte counts) in a Celly 70 auto analyzer (Chronolab, Mexico City, Mexico). Serum was obtained from samples without additives and was tested for glucose, cholesterol, triglycerides and high-density lipoprotein cholesterol (HDL-c) concentrations using a Cobas C111 analyzer (Roche Diagnostics, Basilea, Sw). The intra- and inter-assay variation coefficients were less 5% in all determinations.

After clinical history and physical examination were conducted, we performed the following anthropometric measurements: weight was measured while the woman was wearing underwear and a clinical gown and in a fasting state (after evacuation). A Torino® scale (Tecno Lógica, Mexicana, Mexico, TLM®) was used, and was calibrated before each weight measurement. Height was obtained with an aluminum cursor stadiometer graduated in millimeters. The woman stood barefoot, back, and head in contact with the stadiometer in Frankfurt horizontal plane. Body mass index (BMI) was calculated by dividing weight (in kilograms) through squared height (in meters).

Blood pressure was measured in both arms 3 times in the morning, in a fasting condition, in sitting position. A mercurial manometer was used to measure the blood pressure and it was taken by medical technicians who had attended training sessions to standardize the procedures. The technicians were supervised to avoid possible biases in measurement.

### Assessment of oxidative stress

With the blood samples containing heparin, we measured red blood cell superoxide dismutase (SOD) and glutathione peroxidase (GPx) activities, plasma total antioxidant status (TAS), and plasma malondialdehyde level (MDA). All the methods were validated in our research laboratory, and the within-run precision for the markers were as follows: 3.8%, 4.6%, 4.3%, and 6%. Artefactual formation of thiobarbituric acid reacting substances (TBARS) in the samples was prevented by adding 10 μL of 2 mM butylated hydroxytoluene in ethanol at 95% immediately after centrifugation.

SOD activity was measured by the method that employs xanthine and xanthine oxidase to generate superoxide radicals, which react with 2-(4-iodophenyl)-3-(4-nitrophenol)-5-phenyltetrazolium chloride to form a red formazan dye (Randox Laboratories, Ltd., Crumlin Co. UK). GPx was measured using the oxidation of glutathione by cumene hydroperoxide in the presence of glutathione reductase and NADPH, oxidized glutathione is immediately converted into the reduced form with the subsequent oxidation of NADPH to NADP^+^ (Randox Laboratories, Ltd.). Antioxidant status (TAS) quantification was conducted using 2,2-azino-bis (3-ethylbenzthiazoline-6-sulfonic acid, ABTS^+^) radical formation kinetics (Randox Laboratories Ltd.). The MDA level was measured with TBARS assay, which was performed as described by Jentzsch et al.^11^, and as we previously validated. All the measures were performed in a Shimadzu UV-1601 UV-Vis spectrophotometer (Kyoto, Japan).

Uric acid level was measured by uricase colorimetric method and albumin level by bromocresol green technique with a Cobas C111 analyzer.

In addition, we calculated the antioxidant gap (GAP) with the equation GAP = TAS - [(albumin (μmol) X 0.69) + uric acid (μmol)] [12]

Also, we obtained the SOD/GPx ratio, a proposal from some authors that indicates oxidative damage because the enzymatic antioxidant pathway is a two-step process. In first step, SOD converts superoxide anion to hydrogen peroxide, a strong oxidant; and in the second-step glutathione peroxidase converts hydrogen peroxide to water. Thus, when there is an imbalance between the first and second-step, an accumulation of hydrogen peroxide is produced which affects the cellular functions and may be lead to organic dysfunction [13-15].

Alternative cut-off values of each parameter were defined based on the 90^th^ percentile of young healthy subjects: MDA ≥ 0.320 μmol/L, SOD ≤ 1.20 U/gHb, GPx ≤ 50.1 U/gHb, TAS ≤ 900 μmol/L, SOD/GPx ≥ 0.023, GAP ≤ 190 μmol/L. The uric acid cut-off value was the median of the reference interval (> 268 μmol/L) as determined at the Gerontologic Clinical Research Laboratory of the Universidad Nacional Autónoma de México (UNAM) Zaragoza Campus in Mexico City [10]. An oxidative stress score (SS) was obtained, ranging from 0 to 7, represented the severity of the marker modifications; a score of 1 was given to each value higher or lower than the cut-off point established. A cut-off value of ≥ 4 was considered as OS.

### Assessment of hot flashes, symptomatology linked to menopausal transition and pro-oxidant lifestyle factors

As potential confounding factors were considered mood disturbances, insomnia and pro-oxidant lifestyle aspects. All the women completed Spanish versions of self-assessment tests and a structured questionnaire about pro-oxidant factors.

Menopausal symptoms were assessed with the Menopause Rating Scale (MRS), a validated test to assess the intensity from them [16,17]. The test is composed by 11 items assessing menopausal symptoms divided into three subscales: somatic, psychological and urogenital [18]. Each item can be graded by the subject from 0 (not present) to 4 (1 = mild; 2 = moderate; 3 = severe; 4 = very severe). We used the question about vasomotor symptoms of somatic subscale to assess HFs intensity and strengthen the concepts pictorially.

Anxiety was evaluated with Zung Self-Rating Anxiety Scale (SAS). The SAS is a 20-item measure developed to assess the frequency of anxiety symptoms based on diagnostic conceptualizations. The total scores on the SAS ranged from 0 to 80. A cut-off value >45 was considered to indicate anxiety [19,20].

For depressive mood, we used the Zung Self-Rating Depression Scale (SDS) that consists of 20 items. The score ranges from 20 to 80. A woman with a SDS score below 40 was considered normal [21,22].

We used the Athens Insomnia Scale (AIS) to evaluate sleep disturbances. The AIS is a validated self-assessment psychometric instrument designed to determine sleep difficulty based on the ICD-10 criteria. It consists of eight items and the higher the score, the greater intensity of sleep disturbances. A cut-off value of ≥ 8 was considered as insomnia [23,24].

About lifestyle pro-oxidant factors, the participants answered a structured questionnaire assessing the following: smoking, the consumption of caffeinated and/or alcoholic beverages, and physical inactivity. We considered a pro-oxidant factor present when the following were noted: smoking ≥ 2 cigarettes/day, consumption of ≥ 2 glasses/day alcoholic beverages, consumption of > 2 cups/day caffeinated beverages, and < 30 min/day of physical activity.

### Statistical analysis

Quantitative results were described with the means ± standard error (SE), and they were compared using two sample t-test. We separated the women in three subgroups for each group per the HFs intensity: 1) no/mild (< 2), 2) moderate (= 2), and 3) severe/very severe (≥ 3), and we compared with one-way ANOVA with Dunnett test as *posthoc*, using subgroup 1 as control. Categorical data were analyzed using frequencies, percentages and 95% confidence interval (95%CI) for proportions, which were compared using the chi square test. Also, we calculated Spearman’s correlation between SS and HFs intensity or other tests scores, for each group. Three logistic regression models, with the enter method, were generated according to different confounding factors, using categorical OS (SS cut-off value ≥4) as dependent variable. In all models, we included HFs as no/mild, moderate and severe/very severe, and the other variables as dummy. The first model was unadjusted, only HFs severity was included as independent variable; in the second model, we added anxiety (score > 45), depressive mood (score ≥ 40) and insomnia (score ≥ 8) as confounding pro-oxidant symptoms. Finally, to simultaneously control the risk factors for OS, we incorporated the following variables at the second model: age (≥ 50 y), smoker (> 2 cigarettes/d), caffeinated beverages intake (> 2 cups/d), sedentary (< 30 min/d of physical activity); alcohol intake was not included because the frequency in the groups was very low, and obesity was not added because its frequency was very high. The models were built using the variables identified in the literature as potential pro-oxidant factors associated with OS. Interactions among pro-oxidant variables were not important to the models. With the odds ratio (OR) results, we calculated chi square for trends. Risk factors were defined by OR > 1 and a 95%CI that did not include the 1.0 value. A *p*-value < 0.05 was considered significant. The data were processed using the standard statistical software package SPSS V. 20.0 (IBM SPSS Statistics Armonk, NY, USA).

## Results

### Sample characteristics

A total of 315 women separated in two groups (145 premenopausal and 170 postmenopausal) were included in the study, from a baseline sample of 476 women recruited. Seventy-three (50%) premenopausal women reported mild to moderate hot flashes vs. 112 (66%) postmenopausal women that indicated moderate to severe hot flashes (*p*< 0.01). The biochemical-hematologic parameters, anthropometric and blood pressure measurements in both groups had similar values in all parameters, except in red blood parameters and cholesterol (*p*< 0.0001). Of the analyzed symptoms, insomnia was more frequent in postmenopausal women than in premenopausal women (*p*< 0.05); psychological alterations were not different between the groups. Frequency in the pro-oxidant factors was similar, except that more premenopausal women were smokers (Table 1).

**Table 1.** Descriptive characteristics of study groups.

| Characteristic | Premenopausal women<br>(n = 145) | Postmenopausal women<br>(n = 170) |
| --- | --- | --- |
| Age (y) | 47.1 ± 0.3 | 52.9 ± 0.3<sup>a</sup> |
| Estrogen (pg/mL) | 94.3 ± 6.3 | 10.1 ± 0.5<sup>a</sup> |
| FSH (mU/mL) | 17.0 ± 1.6 | 57.5 ± 2.0<sup>a</sup> |
| Hemoglobin (mmol/L) | 8.3 ± 0.07 | 9.0 ± 0.06<sup>a</sup> |
| Hematocrit (%) | 42.8 ± 0.29 | 43.9 ± 0.30<sup>b</sup> |
| Glucose (mmol/L) | 5.27 ± 0.13 | 5.38 ± 0.13 |
| Cholesterol (mmol/L) | 5.28 ± 0.08 | 5.77 ± 0.09<sup>a</sup> |
| Triglyceride (mmol/L) | 2.09 ± 0.12 | 2.15 ± 0.11 |
| HDL-c (mmol/L) | 1.42 ± 0.03 | 1.48 ± 0.03 |
| Systolic tension (mm Hg) | 122 ± 1.4 | 125 ± 1.2 |
| Diastolic tension (mm Hg) | 82 ± 0.9 | 84 ± 0.7 |
| Body mass index (kg/m<sup>2</sup>) | 29.72 ± 0.7 | 29.52 ± 0.4 |
| Anxiety | 38 (26%, 19-33%) | 55 (32%, 25-39%) |
| Depressive mood | 35 (24%, 17-31%) | 51 (30%, 23-37%) |
| Insomnia | 74 (51%, 43-59%) | 110 (65%, 58-72%)<sup>c</sup> |
| Smokers (> 2 cigarettes/d) | 28 (19%, 13-25%) | 15 (9%, 5-13%)<sup>c</sup> |
| Caffeinated beverages intake (> 2 cups/d) | 49 (34%, 26-42%) | 48 (28%, 21-35%) |
| Alcohol intake (> 2 glasses/d) | 7 (5%, 1-9%) | 7 (4%, 1-7%) |
| Sedentary (<30 min/d of physical activity) | 90 (62%, 58-66%) | 99 (58%, 51-65%) |

Quantitative data show means ± standard error; categorical data show frequency, percentage and 95% confidence interval. Two sample t-test, <sup>a</sup>*p* < 0.0001, <sup>b</sup>*p* = 0.01; <sup>c</sup>chi square test, *p* < 0.05. FSH: Follicle stimulating hormone, HDL-c: high-density lipoprotein cholesterol.

### Oxidative stress and hot flashes

Among OS markers, MDA level was higher in postmenopausal women with severe HFs compared to women with mild HFs (*p*< 0.01), and SOD activity was lower (*p*< 0.05) when HFs intensity increase in this group. In premenopausal women, the markers did not show any change with HFs severity. Additionally, we used an oxidative stress score (SS) that integrates both oxidized and antioxidant markers to represent the dynamics of OS. This index included MDA level and SOD/GPx ratio as oxidative damage markers, two antioxidant enzymes (SOD and GPx), and three plasma antioxidant components (TAS, GAP and uric acid), this to evaluate integrally the OS. In this context, we found that SS was increased with HFs severity (mild 2.9 ± 0.23, moderate 3.1 ± 0.21 and severe 3.8 ± 0.18, p<0.01) in postmenopausal women; in premenopausal women, the index did not change (Table 2).

**Table 2.**
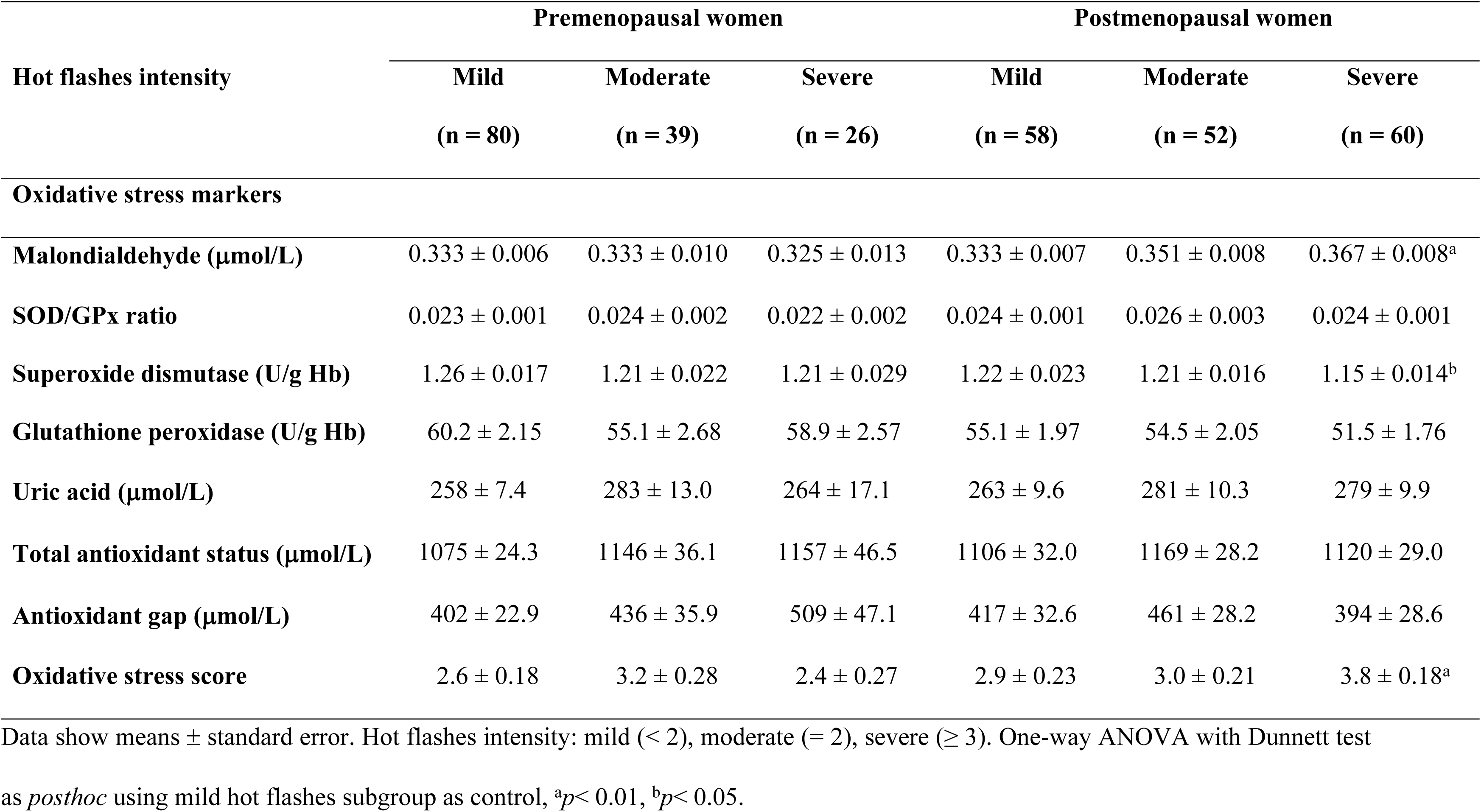
Oxidative stress markers by host flashes intensity in study groups.

In the univariate analyses between SS and HFs severity or each menopausal symptom measured by the applied test scores, we found a better correlation between SS and HFs intensity (r = 0.247, p= 0.001) in postmenopausal women; other menopausal symptoms were not significant. In premenopausal women, we did not see any relationship (Table 3).

**Table 3.** Relationship between stress score and menopausal symptoms scores in study groups.

| Scale | Premenopausal women<br>(n = 145) |  | Postmenopausal women<br>(n = 170) |  |
| --- | --- | --- | --- | --- |
|  | r | p value<sup>a</sup> | r | p value<sup>a</sup> |
| **Hot flashes intensity** | 0.087 | 0.298 | 0.247 | 0.001 |
| **MRS score** | 0.029 | 0.730 | 0.148 | 0.054 |
| **AIS score** | 0.052 | 0.543 | 0.062 | 0.431 |
| **SAS score** | 0.117 | 0.169 | 0.059 | 0.457 |
| **SDS score** | 0.045 | 0.598 | 0.053 | 0.508 |

<sup>a</sup>Spearman correlation. SDS: Zung Self-Rating Depression Scale; MRS: Menopause Rating Scale; AIS: Athens Insomnia Scale; SAS: Zung Self-Rating Anxiety Scale.

Furthermore, we built several logistic models that included HFs intensity, menopausal symptoms and lifestyle pro-oxidant factors. We found significant models for postmenopausal women (Table 4). Accordingly, we observed a gradual increment of risk for OS in postmenopausal women with severe HFs when we included pro-oxidant factors; thus, when the model was unadjusted, OR = 2.54 (95% CI: 1.20-5.39), and when we incorporated both, menopausal symptoms and lifestyle pro-oxidant factors as categorical variables, the risk increased to 3.37 (95% CI: 1.20-9.51)

**Table 4.** Odds ratio to present different intensities of hot flashes according to oxidative stress and pro-oxidant factors in postmenopausal women.

| Hot flashes intensity<sup>a</sup> | OR (95% confidence interval) |  |  |
| --- | --- | --- | --- |
|  | Mild (< 2)<br>(n = 58) | Moderate (= 2)<br>(n = 52) | Severe (≥ 3)<br>(n = 60) |
| **Oxidative stress (SS ≥ 4)** | 18 (31%) | 18 (35%) | 32 (53%) |
| **Model<sup>b</sup>** |  |  |  |
| **A** | 1.00 | 1.18 (0.53-2.61) | 2.54 (1.20-5.39)<sup>c</sup> |
| **B** | 1.00 | 1.62 (0.69-3.83) | 3.33 (1.35-8.21)<sup>d</sup> |
| **C** | 1.00 | 1.51 (0.57-3.95) | 3.37 (1.20-9.51)<sup>e</sup> |

SS: stress score.
<sup>a</sup> According MRS scale.
<sup>b</sup> Models included following baseline variables:
A. Unadjusted model included hot flashes severity and oxidative stress score ≥ 4. Significance of the model $p=0.03$ ; <sup>c</sup> chi square for trend $p=0.01$ .
B. Adjusted for: anxiety (score > 45), depressive mood (score ≥ 40) and insomnia (score ≥ 8). Neither of the symptoms were significant for the model. <sup>d</sup> Significance into the model $p<0.01$ .
C. Add to B model the pro-oxidant factors: age (≥ 50 y), smoker (> 2 cigarettes/d), caffeinated beverages intake (> 2 cups/d), and sedentary (< 30 min/d of physical activity). Neither of the pro-oxidant factors were significant for the model. <sup>e</sup> Significance into the model $p<0.05$ .

## Discussion

Hot flashes are the most prevalent and bothersome symptoms reported by women during the menopausal transition; recently it was noted that up to 80% of women experience HFs during this period and that on average, symptoms persist at least 5 years [4]. HFs intensity increase around perimenopause and is high in the postmenopausal period [4,25]. As we observed in this study, postmenopausal women referred more discomfort due to HFs because their vasomotor sensations were moderate to severe, contrary to premenopausal women; however, the prevalence of moderate/severe HFs in our study was higher than the Penn Ovarian Aging Study cohort [4] and other study [26] (66% vs. 46% and 40%), this difference may be due to the cross-sectional design of our study and the way of collecting the data; moreover, the populations are different and the HFs are dependent on several factors such as genetic, diet, physical changes, cultural influences, and individual experiences and expectations [27].

Furthermore, HFs have been consistently shown to be associated with discomfort, sleep disturbances, fatigue, mood disturbances and deficient quality of life [25,28], all pro-oxidant factors that cause OS. Oxidative stress occurs when the balance between ROS, produced by the metabolism, and antioxidants is disrupted, causing an accumulation of reactive species or the depletion of antioxidants [29,30]; this imbalance can cause severe oxidative damage in cells, and it is related to several chronic diseases that frequently are associated to aging, as well as HFs have been linked to cardiovascular disease, osteoporosis and cognitive decline [27]. In fact, vasomotor symptoms are associated with an increase in carotid intima–media thickness and other vascular changes, which causes vascular dysfunction and activation of pathways that increase the production of ROS and promote OS [31-33].

Additionally, OS is increased in the postmenopausal period probably due to the decrease in estrogen level, a natural antioxidant, by different biochemical mechanisms. Moreover, we previously noted that menopause is a risk factor for OS, which may be due to an estrogenic deficiency and symptomatology severity [8]; thus, in this study we explored which of the symptoms may be the cause of OS, and we focused on HFs because it is the onset of all disturbances.

In this sense, we observed that MDA level was higher and SOD activity was lower, as individual markers, and SS was higher, all in postmenopausal women with severe HFs, showing high OS in these women. Although there are few references that analyze the relationship between HFs and OS, our results are similar to a study in which postmenopausal women with HFs had a high lipoperoxide level (MDA) and low total antioxidant status compared to women without HFs [34]; and other research that showed a markedly reduced antioxidant defense in women with vasomotor symptoms [35], probably because E2 can stimulate cellular antioxidant enzymes [6,36], and this capacity is lost in the postmenopausal period. However, recently a research was conducted to evaluate the association between HFs and OS markers in middle-aged women; this study indicated that none of the peripheral markers examined were found to be significantly associated to the presence of HFs [9], contrary to our results. A possible explanation to these controversial results is that the authors used urinary 8-iso-prostaglandin F2α and 8-OH-deoxy-2’-guanosine level as oxidative damage markers and different antioxidant components, and in our study, we used an index to integrally assess OS that includes both, oxidized and antioxidant components. Although some oxidative damage markers are used to evaluate OS, such as 4-HNE and isoprostanes, the measurement of antioxidant markers has not always been consistent, therefore, our research group had proposed an index to integrate both processes. This index has been used in different studies and shown to be useful for measuring OS [37-39].

Besides, it is known that the psycho-neuro-endocrine change and vasomotor symptoms during menopausal transition affect self-esteem, mood states and therefore quality of life [40,41]. Furthermore, depressive mood, anxiety and insomnia are considered pro-oxidant factors as well as the severe discomfort felt, and these factors can increase OS [8,39,42,43]. Therefore, we analyze a possible correlation between SS and menopausal symptoms scores to assess which alteration was related with OS. We found a positive correlation with HFs severity in postmenopausal women, but the other symptoms tested did not show any relationship; and these women with severe HFs had two-fold more risk for OS compared to women with mild HFs, even after controlling for menopausal symptoms and pro-oxidant factors. In this sense, although there are inconsistencies in the reports about the relationship between HFs and OS, our results showed that postmenopausal women with severe HFs have a high risk for OS.

In the physiology of HFs, the sweat and vasodilation produced by the process are controlled by the thermoregulatory nucleus, located in the preoptic area of the hypothalamus, which regulates core body temperature to maintain a homeostatic range (thermoregulatory zone). This thermoregulatory zone is narrow in postmenopausal women; therefore, small increases in core body temperature can trigger HFs [3,44]. The thermoregulatory zone is controlled by a complex neuroendocrine pathway that can produce an increment of norepinephrine and serotonin that causes changes in the thermoregulatory nucleus, diminishing the set point and increasing the probability of HFs [45]. The rapid degradation of these monoamine neurotransmitters is fundamental for the correct synaptic neurotransmission, but it is also a reaction that involves the enzyme monoamine oxidase (MAO), which generates several products, such as hydrogen peroxide, that are potentially neurotoxic and can trigger the production of ROS and induce mitochondrial damage and neuronal apoptosis, increasing the OS [46]. Additionally, in the setting of OS, catecholamines are oxidatively converted to different molecules that are potentially oxidants, which may further develop an environment of OS. An OS environment leads eventually to cytotoxic responses and altered cellular function, producing neuronal degeneration [47]. Thus, the women with severe HFs are in a constant oxidative challenge without the protection of estrogen as the main antioxidant mechanism at both cerebral and systemic level, since the women have estrogen receptors in many cells of the body [6], which probably increases their OS.

It is necessary to consider several limitations of this study. Initially, we do not include some pro-oxidant factors such as certain types of food in the diet and the cooking style, but all the participants had similar lifestyles and socioeconomic level, therefore we supposed similar feeding habits; besides we considered the main pro-oxidant factors in the analysis. Other limitations are study design, which can be used to explore the associations between HFs and OS but is unable to establish a causal conclusion, and the sample size. Even so, the sample size that allowed us to achieve similar subgroups size when we stratified by HFs severity in postmenopausal women, the use of an integral OS index and the control of potential confounders in the multivariate logistic models, are aspects that strengthen the study.

In conclusion, among Mexican postmenopausal women there is an association between HFs severity and OS; however, longitudinal studies or controlled clinical trials must be carried out to confirm our findings.

## Author contributions

MASR conceived the study, completed statistical analyses and interpretation of data, and drafted the manuscript. MZF conceived the study and contributed to the interpretation of data. AAR collected and interpreted the data. VMMN contributed to the discussion and revised the manuscript. All authors read and approved the final manuscript.

